# Natural transformation protein ComFA exhibits single-stranded DNA translocase activity

**DOI:** 10.1101/2021.10.29.466550

**Authors:** Hannah R. Foster, Xiaoxuan Lin, Sriram Srikant, Rachel R. Cueny, Tanya G. Falbel, James L. Keck, Rachelle Gaudet, Briana M. Burton

## Abstract

Natural transformation is one of the major mechanisms of horizontal gene transfer in bacterial populations and has been demonstrated in numerous species of bacteria. Despite the prevalence of natural transformation, much of the molecular mechanism remains unexplored. One major outstanding question is how the cell powers DNA import, which is rapid and highly processive. ComFA is one of a handful of proteins required for natural transformation in gram-positive bacteria. Its structural resemblance to the DEADbox helicase family has led to a long-held hypothesis that ComFA acts as a motor to help drive DNA import into the cytosol. Here, we explored the helicase and translocase activity of ComFA to address this hypothesis. We followed the DNA-dependent ATPase activity of ComFA and, combined with mathematical modeling, demonstrated that ComFA likely translocates on single-stranded DNA from 5’ to 3’. However, this translocase activity does not lead to DNA unwinding in the conditions we tested. Further, we analyzed the ATPase cycle of ComFA and found that ATP hydrolysis stimulates the release of DNA, providing a potential mechanism for translocation. These findings help define the molecular contribution of ComFA to natural transformation and support the conclusion that ComFA plays a key role in powering DNA uptake.

**Importance:** Competence, or the ability of bacteria to take up and incorporate foreign DNA in a process called natural transformation is common in the bacterial kingdom, but understanding of the mechanism is still limited. Increasing evidence in several bacteria confirms that long, contiguous stretches of DNA are imported into cells, and yet how bacteria power processive transformation remains unclear. Our finding that ComFA, a DExD-box helicase required for competence in gram-positive bacteria, translocates on single-stranded DNA from 5’ to 3’, supports the long held hypothesis that ComFA may be the motor powering DNA transport during natural transformation. Moreover, ComFA may be a previously unidentified type of DExD-box helicase—one with the capability of extended translocation on single-stranded DNA.

## Introduction

Natural transformation, or the uptake and integration of extracellular DNA by competent bacteria, is one of the main mechanisms for sharing genetic information among bacterial populations (1,2). Natural transformation is unique among the methods of horizontal gene transfer because the process is entirely controlled by the DNA recipient—the DNA donor organism need not be present (3). Competence is relatively common in the bacterial kingdom, having been demonstrated in at least 80 species, and the competence proteins and basic mechanism of transformation have been defined in both gram-positive and gram-negative bacteria (3–5). However, key aspects of the molecular mechanism of DNA uptake remain a mystery.

One such mystery is the source of power for DNA transport across the cell membrane. The speed and processivity of DNA uptake strongly suggest that the cell uses energy to actively transport DNA. Indeed, magnetic tweezer experiments that tracked the movement of *Bacillus subtilis* along a DNA substrate demonstrated that uncoupling agents could halt the DNA uptake process, confirming that the process is dependent in some way upon the consumption of ATP (6). One hypothesis is that a DNA uptake motor helps power the translocation of DNA. However, a motor coupling ATP hydrolysis to DNA uptake has yet to be identified.

The process of natural transformation begins with the binding of double-stranded DNA (dsDNA) outside the cell (7,8). Gram-positive and gram-negative bacteria differ in the early steps to bring dsDNA across the cell wall and outer membrane (if present), but then in all naturally competent bacteria, ComEA guides the dsDNA toward a pore in the cell membrane (7,8). At this point, a single strand of DNA is taken up into the cell, while the other is degraded (9–11). ComEC is presumed to be the channel protein that allows for the passage of DNA through the cell membrane, based on its large size, its conservation in every competent species identified to date, and the fact that it is indispensable for transformation (3,12,13). In gram-positive bacteria, membrane-associated ComFA is also necessary for the transport of DNA across the cell membrane (9–11,14,15). The precise role of ComFA has not been identified, but researchers have put forth the hypothesis that the protein could act as a DNA uptake motor in gram-positive bacteria (1,16–19)

The idea that ComFA may be the DNA uptake motor, in part, stems from its importance in natural transformation. Transformation efficiency is decreased by as much as 10,000-fold in *B. subtilis* strains lacking ComFA (16–18). Similarly, competence in *Streptococcus pneumoniae* strains lacking ComFA is dramatically reduced or eliminated (20,21). In addition to its critical role in transformation, however, the structural similarity of ComFA to the DEAD-box family of helicases, which belongs to helicase superfamily 2, led to the hypothesis that ComFA may push or pull DNA into the cell. ComFA contains a DEAD-box helicase-like domain, including conserved Walker A and Walker B motifs (17,18). Mutation of either of these domains in *B. subtilis* ComFA significantly reduces transformation efficiency, while having no effect on DNA binding to the cell surface (17,18). Our previous findings that *S. pneumoniae* ComFA binds single-stranded DNA (ssDNA) and that the protein exhibits ssDNA-dependent ATPase activity bolster the hypothesis that ComFA is the DNA uptake motor (19). However, DEAD-box helicases are known principally as RNA remodelers that bind and melt nucleic acids locally, rather than unwinding DNA via translocation (22–24). This raises the question, by what mechanism could ComFA couple ATP hydrolysis with DNA uptake?

Here, we present biochemical data supporting the model that ComFA acts as a DNA translocase in gram-positive bacteria, making it unique among DEAD-box helicases. We use the ComFA protein from *S. pneumoniae* to demonstrate that ComFA does not bind dsDNA and that it does not require a free 5’ or 3’ end to bind ssDNA. ATPase assays and mathematical modeling reveal that ComFA, unlike most DEAD-box helicase family proteins, likely translocates on DNA in a 5’ to 3’ direction. We used a strand displacement assay to determine that the translocase activity of ComFA does not give it the ability to unwind DNA under the conditions we tested. Further, we analyzed the coupling of the ATPase cycle to ssDNA interaction. Based on these data, we propose a mechanism for DNA translocation of ComFA based on ATP and ADP binding dynamics.

## Results

### ComFA binds dsDNA very weakly

We showed previously that ComFA ATPase activity is not stimulated by doublestranded DNA (dsDNA; 19). However, to further elucidate the role of ComFA in DNA uptake, we sought to determine whether ComFA is capable of binding dsDNA. We tested the dsDNA-binding ability of ComFA using an Electrophoretic Mobility Shift Assay (EMSA) and found that ComFA associates with dsDNA only very weakly, with a K_d_ of >1 μM, as opposed to 138 ± 86 nM for ssDNA which we reported previously (Figure 1A-B; 19). To further test the preference for ssDNA, we performed a competition experiment, examining the previously demonstrated ssDNA-stimulated ATPase activity of ComFA (19) in the presence of increasing concentrations of dsDNA. We measured the ATPase activity of ComFA while keeping the concentration of ssDNA constant at 500 nM and increasing the concentration of dsDNA to see whether high concentrations of dsDNA could compete away the interaction of ComFA with ssDNA (Figure 1C). Presence of dsDNA failed to decrease the ATPase activity of ComFA even at a concentration 20 times that of the ssDNA, confirming that ComFA exhibits a strong preference for ssDNA.

**Figure 1.**
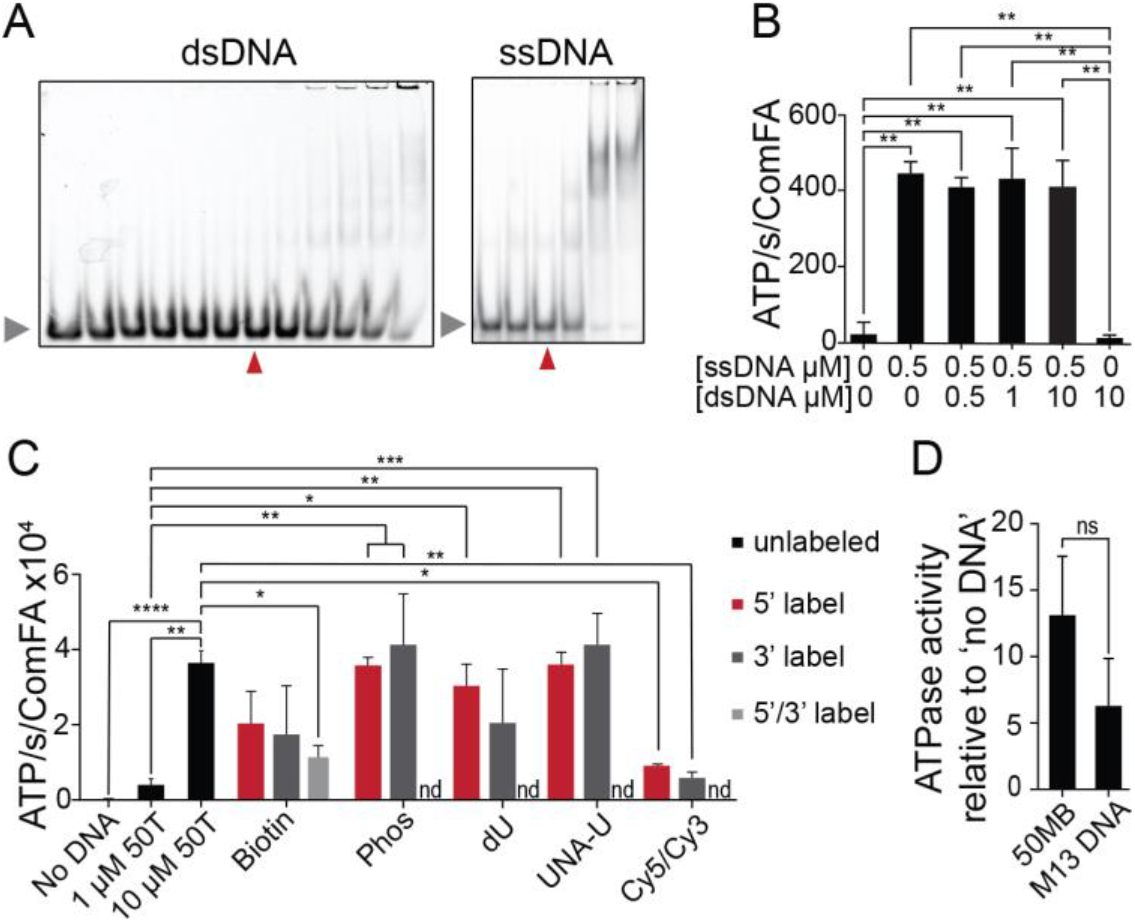
ComFA binds weakly to dsDNA and binds internally on ssDNA. (A) A representative Electrophoretic Mobility Shift Assay (EMSA) using 5 nM dsDNA with increasing concentrations of ComFA (dsDNA gel on the left: 0, 10, 20, 40, 60, 80, 100, 200, 400, 600, 800, 1200 nM; ssDNA gel on the right: 0, 50, 100, 200, 400, 800 nM); red triangle indicates 100 nM). dsDNA was made with 50MB and 50MB-rev (see Table 1 for sequences). ssDNA oligo was 50MB. Contrast and brightness were adjusted slightly for presentation. (B-D) Mean ATPase activity of WT ComFA with indicated substrates. ATPase activity was evaluated using an NADH-coupled 96-well plate assay with and without nucleic acid substrate as indicated (concentration in μM) (B) n=2 all conditions. ssDNA=50T, dsDNA=50MB/50MB-rev. (C) No DNA, 1 or 10 μM 50T, and 10 μM 50T labeled with biotin (single-labed; n=3 or dual-labeled 5’/3’; n=2), phosphorylation (Phos; n=2), deoxyuridine (dU; n=3), UNA-U (n=3), fluorescent tag (Cy5 or Cy3; n=2). (D) [DNA]=250μg/ml. Due to high apparent background ATPase activity of two replicates, these data are shown relative to the ‘no DNA’ control. Error bars represent standard deviation. N=4. (B-C) A one-way ANOVA with Sidak’s multiple comparisons test was used and for (D) an unpaired t-test was used to determine significance, denoted by *p≤0.05, **p≤0.01, *** p≤0.001,****p≤0.0001.

### ComFA binding does not require a free end of DNA

Many ssDNA binding proteins bind or load exclusively on either the 5’ or 3’ end of their respective DNA substrates, and this binding preference can be critical for the role they play in the cell (28). Determining whether ComFA requires a free DNA end could shed light on when during the uptake process ComFA interacts with DNA. To this end, we tested the ATPase activity of ComFA with dT_50_ oligonucleotides (50T) that were blocked on either the 3’ or the 5’ end or both ends by modifications, such as biotin, phosphorylation (Phos), deoxyuridine (dU), UNA-U, or fluorescent tags (Cy5/Cy3). We reasoned that if ComFA requires a free end of DNA, then blocking that end with a modification should decrease ATPase activity. Most modifications did not affect the stimulated ATPase activity. Oligos modified by a fluorescent tag did induce a slightly lower (<7-fold) ATPase activity than unmodified oligos, as did oligos dual-labeled with biotin. However, the differences were not end-specific and therefore likely arose due to steric effects created by the larger tags (Figure 1D).

To further test whether a free end of DNA was not essential for ComFA binding, we measured ComFA stimulation by single-stranded, circular M13mp18 bacteriophage DNA. Since M13mp18 DNA is much longer than ssDNA oligos, we used the equivalent number of bases of either M13mp18 DNA or 50MB; we added 250 μg/ml of either M13mp18 DNA or ssDNA oligo. Although M13mp18 DNA may not stimulate ATPase activity to the degree of single-stranded oligos, this difference was not significant and may be attributed to the fact that M13mp18 DNA is more likely to fold into complex secondary structures than the single-stranded oligo we designed specifically to limit secondary structure (Figure 1E). These data lead us to conclude that ComFA does not require an unobstructed end of DNA and therefore is capable of binding internally on DNA.

### ComFA ATPase activity is DNA length-dependent

Helicases that translocate on nucleic acid substrate frequently show a dependence on substrate length (26,27). That is, longer substrates stimulate higher ATPase activity, since longer substrates allow for further ATP-stimulated “travel” when the proteins are sufficiently processive. We examined the ATPase activity of ComFA in response to increasing lengths of ssDNA (5T to 100T) to determine whether ComFA might translocate on its substrate. Lengths of DNA less than or equal to 20 bases failed to significantly stimulate ATPase activity (Figure 2A; Supplemental Figure S1A-C), but with longer substrates up to 80 bases in length, ComFA exhibited ATPase activity that increased with DNA length (Figure 2A). Even over a range of substrate concentrations from 1-100 μM, longer oligos consistently stimulated higher ATPase activity (Figure 2B, 2C).

**Figure 2.**
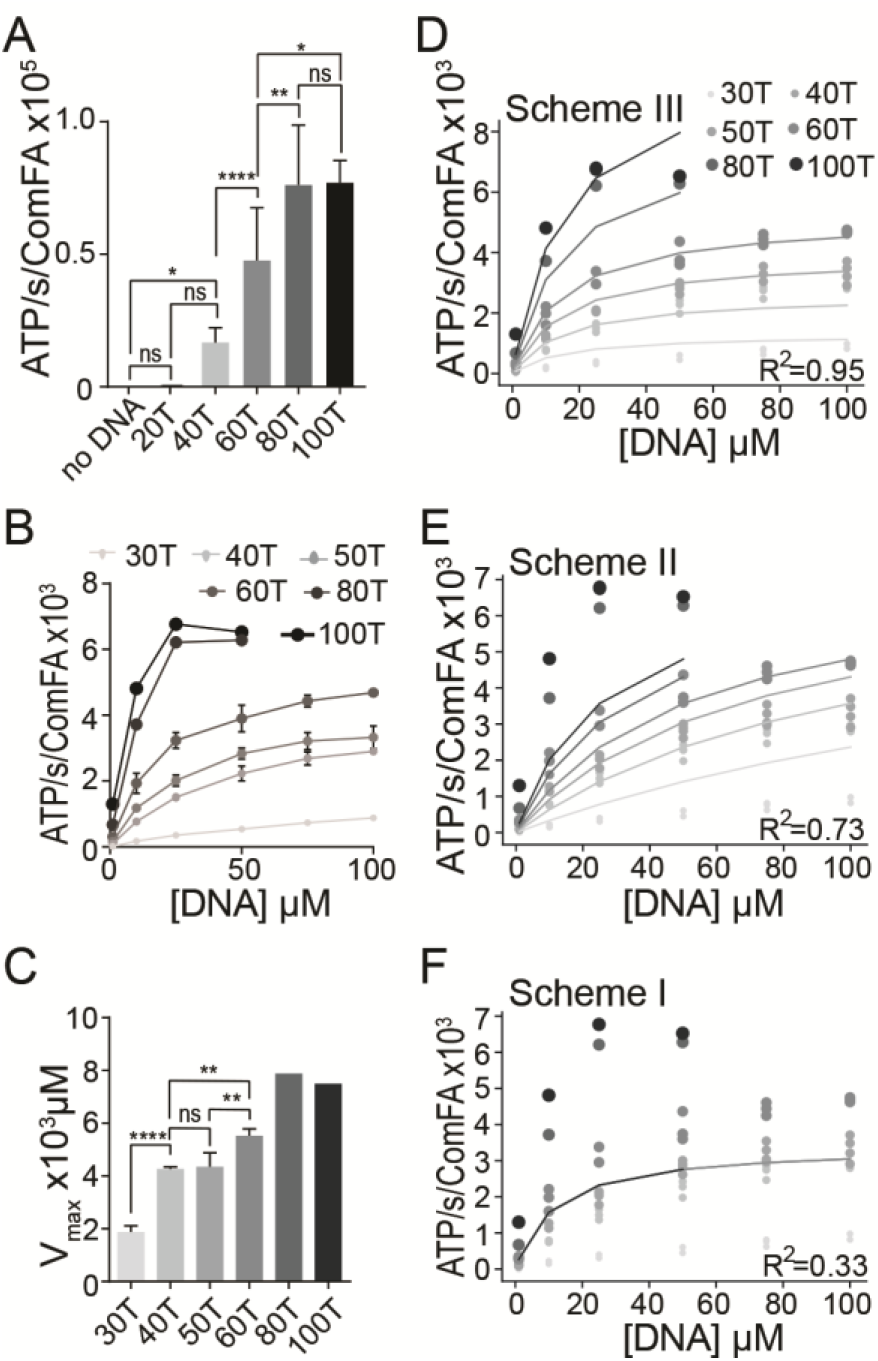
ATPase activity of ComFA is dependent on DNA length. (A) Mean ATPase activity of WT ComFA with poly-dT substrate of various lengths at a concentration of 10 μM. no DNA n=20, 20T n=2, 40T n=7, 60T n=7, 80T n=5, 100T n=4 (B) Mean ATPase activity of ComFA with poly-dT substrate of various lengths and concentrations. ATPase activity was evaluated using an NADH-coupled 96-well plate assay with indicated concentration and length of poly-dT ssDNA. 40T n=3, 50T n=4, 60T n=3, 80T n=1, 100T n=1. (C) Mean Vmax (1-100 μM) for indicated lengths of DNA from the data in (B). (D-F) ComFA ATPase activity plotted as dots alongside Scheme III (D), Scheme II (E), or Scheme I (F) of the model, shown as lines of various shades of gray. Length of DNA is indicated by dot size and shade of gray, with lighter shades of gray and smaller dots indicating shorter lengths of DNA. Error bars represent standard deviation. For (A) and (C), a one-way ANOVA with Tukey’s multiple comparisons test was used to determine significance, denoted by *p≤0.05, **p≤0.01, ***p≤0.001, ****p≤0.0001. (D-F) An F-test was used to determine significance of data fit to Scheme III or II models relative to the null model of Scheme I. Scheme III vs Scheme I, F-statistic=335.9; p<0.0001 and Scheme II vs Scheme I, F-statistic=43.0; p<0.0001.

To rule out the possibility that the increase in ATPase activity stimulated by longer oligos is simply due to the increase in the number of nucleotide residues added, we plotted the ATPase activity of ComFA in response to different lengths of oligos against the DNA concentration in μM times the number of nucleotides (μM*nt). If the DNA length is indeed critical, rather than the total number of nucleotides, we would expect, for instance, that the addition of 100T at 25 μM would stimulate a higher ATPase rate than 50T at 50 μM, even though both yield 2500 μM of nucleotide bases. This is, in fact, what we observe: longer oligos stimulate higher ATPase rates even when the number of nucleotides is taken into consideration (Supplemental Figure S2).

Helicase dependence on substrate length has previously been exploited to distinguish between translocation and non-translocation mechanisms (26,27,29,30). In fact, mathematical models have been developed for ATPase data generated from non-translocating and translocating proteins, such that we were able to fit the ATPase data from ComFA to these models. In brief, we fit our data to three different models, based on those previously presented (26,27). Scheme I (null model) represents a non-translocating protein, with dependence on DNA concentration, but not DNA length. Scheme II represents a simple translocating protein that has the same dissociation rate constant in the middle of the substrate as at the end of the substrate. Scheme III represents a complex translocating protein that has two different dissociation rate constants—dissociation within the substrate molecule and dissociation at the end of the molecule (26,27).

The model requires the input of a minimum DNA binding length. While data from EMSAs suggested that ComFA had weaker binding to 30-base sequences (Supplemental Figure S1C), 30-base sequences still stimulated measurable ATPase activity that was significantly higher than the ‘no DNA’ control (Figure 2A; Supplemental Figure S1A-B). It is probable that the steric hindrance we observed with the fluorescently-labeled probe in Figure 1D interfered with the binding of ComFA to smaller fluorescently-labeled substrates. Therefore, we assigned the minimum binding length of ComFA to be 20 bases, such that the model considered ATPase data stimulated by oligos longer than 20 bases.

Scheme III provided the best fit for the ATPase data of ComFA (Figure 2D; R^2^ = 0.95), followed by Scheme II (Figure 2E; R^2^ = 0.73). On the other hand, fitting the data to Scheme I yielded the worst fit (Figure 2F; R^2^ = 0.33). The F-test finds that the data fit either of the translocating models (Scheme III or Scheme II) significantly better than the non-translocating model (null model; Scheme I; p<0.0001), suggesting that the ATPase activity of ComFA in response to oligos of different lengths more closely resembles that of a translocating protein than a non-translocating protein. In addition, the closer fit of our data to Scheme III indicates that ComFA may have two different dissociation constants depending on whether it is dissociating in the middle or at the end of the DNA molecule.

The fact that increasing length past 80 bases did not increase ATPase activity (Figure 2A-B), suggests that ComFA has limited processivity. Because ComFA binds internally on DNA and therefore will, on average, bind in the middle of a strand of DNA, we can conclude that ComFA can translocate up to 30-40 bases before dissociation.

### ATPase activity of ComFA suggests 5’ to 3’ directionality of translocation

Proteins that translocate on single-stranded nucleic acid often exhibit a preferred directionality (i.e. 5’ to 3’ or 3’ to 5’; Singleton et al., 2007; Pyle, 2008). While testing the enzymatic activity of ComFA in response to different nucleic acid substrates, we discovered an experimentally useful trait of ComFA: ComFA reproducibly exhibits roughly 3 times greater ATPase activity with polyT substrates than with mixed-base substrates (Figure 3B). These findings, combined with the data demonstrating that ComFA binds internally to DNA, provided the means to test directionality of translocation. We thus designed two different oligos: an oligo with 30 dTs on the 5’ end and 20 mixed bases on its 3’ end (30T20MB) and an oligo with 20 mixed bases on its 5’ end and 30 dTs on the 3’ end (20MB30T; Figure 3A). If ComFA travels from 3’ to 5’, 30T20MB will stimulate higher ATPase activity, as ComFA will bind internally on the DNA and therefore, on average encounter more polyT than mixed-base sequence (see model in Figure 3A). On the other hand, if ComFA travels from 5’ to 3’, 20MB30T will stimulate higher ATPase activity. We found that ComFA ATPase activity was stimulated to a significantly greater degree by 20MB30T (which stimulated ATPase activity equivalent to 50T) than by 30T20MB or by a 50 mixed-base oligo (50MB; Figure 3B), consistent with the interpretation that ComFA likely translocates from 5’ to 3’.

**Figure 3.**
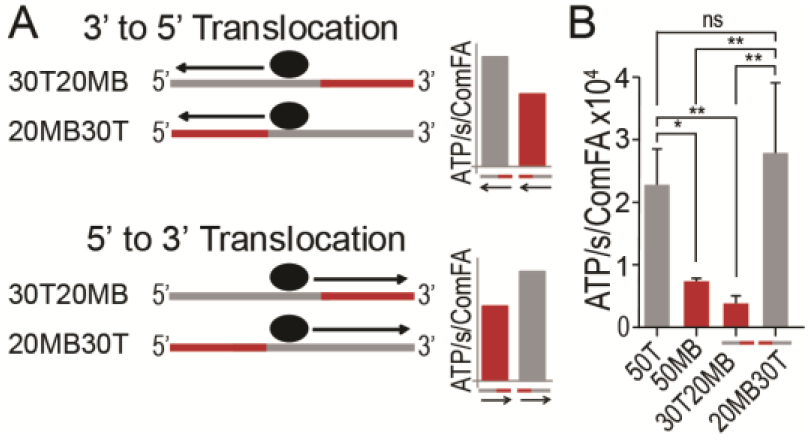
ComFA exhibits 5’ to 3’ directionality. (A) Model of expected outcomes with respective oligos if ComFA travels in a 3’ to 5’ or 5’ to 3’ direction. Gray line indicates poly-dT sequence, while the red line indicates mixed base sequence. Black oval represents ComFA. (B) Mean ATPase activity of ComFA in the presence of indicated oligos (for DNA sequences, see table 1). 50T n=4, 50MB n=3, 30T20MB n=4, 20MB30T n=4. Error bars represent standard deviation. A one-way ANOVA with Tukey’s multiple comparisons test was used to determine significance, denoted by *p≤0.05, **p≤0.01.

### ComFA lacks helicase activity

We next examined ComFA DNA binding and unwinding activities to determine whether ComFA has helicase activity. First, we tested the binding of ComFA without ATP to 50 and 30 polyT sequences (50T, 30T) and to two annealed double/single-stranded oligo combinations, one containing a 5’ dsDNA sequence 20 base pairs in length with a 30 base 3’ dT overhang (5’ ds), and the other with a 20 base pair 3’ dsDNA sequence with a 30 base 5’ dT overhang (3’ ds; Figure 4A; see Table 1 for sequences). ComFA bound either the 5’ ds complex or the 3’ ds complex with equal affinity and with similar affinity as it did 50T or 30T (< 40 nM; Figure 4B). The Kd of 138 ± 86 nM, reported in Diallo et al. (19) was higher than what we report here, but we attribute this to the use of the altered protein purification protocol combined with optimization of the EMSA conditions for these experiments.

**Figure 4.**
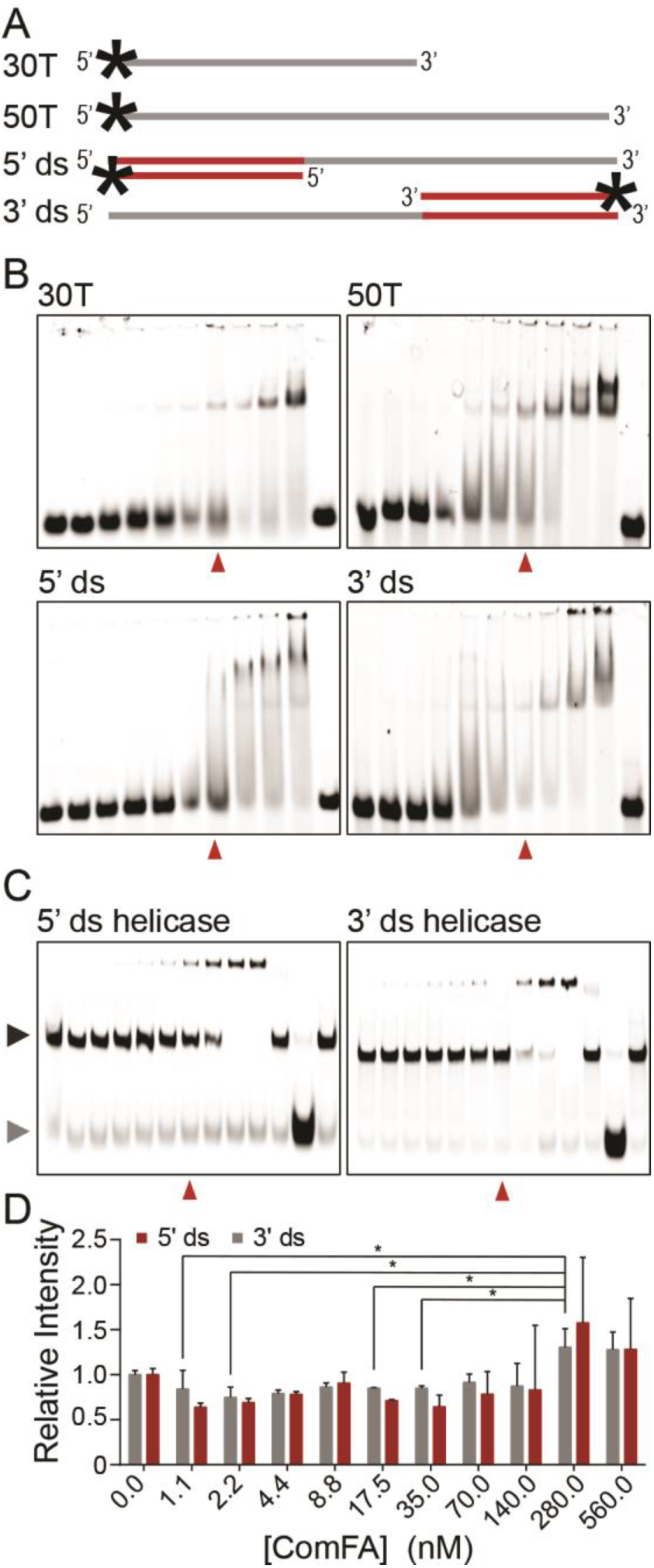
ComFA lacks helicase activity. (A) Diagram of single-stranded and double-stranded oligos used. Red lines indicate mixed base sequences, gray lines indicate poly dT sequences. 3’ ds=30T20MB/20MB annealed, 5’ ds=20MB30T-B/20MB. (B) Representative EMSAs with substrates (40 nM DNA, 1.1, 2.2, 4.4, 8.8, 17.5, 35, 70, 140, 280, 560, 0 nM ComFA). Red triangles indicate 70 nM ComFA. (C) Representative DNA gels from strand displacement assay (40 nM DNA, lanes are: 1.1, 2.2, 4.4, 8.8, 17.5, 35, 70, 140, 280, 560, 0 nM ComFA, 560 nM ComFA without ATP, unfolded substrate, and folded substrate without ComFA). Red triangles indicate 70 nM ComFA. Gray triangle indicates unfolded DNA, black triangle indicates folded (ds) DNA. Contrast and brightness were adjusted slightly for presentation. (D) Quantification (Fig. 4, cont.) of unwinding activity (intensity of unfolded band relative to ‘no ComFA’). Error bars represent standard deviation (n=3).

**Table 1.**
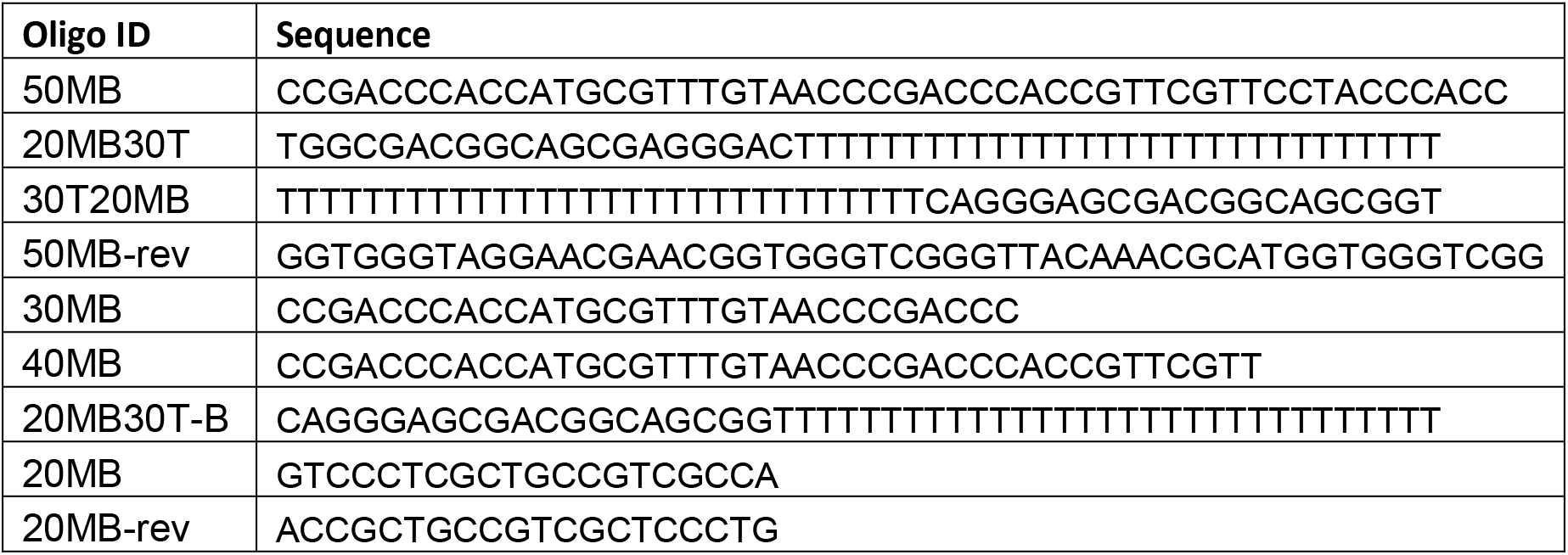
Oligos used in this study. Oligos were ordered from Sigma Aldrich or IDT.

We next used a strand displacement assay to test for DNA unwinding ability of ComFA. We incubated varying concentrations of ComFA with the ss-dsDNA complexes in the presence of ATP to allow strand displacement to occur. We found that ComFA does not unwind dsDNA in our experimental conditions (Figure 4C, D). While the amount of unwound DNA trended up very slightly with the highest concentrations of ComFA, no difference between the 5’ ds and 3’ ds complexes could be detected and the slightly higher values for 280 nM and 560 nM ComFA were not significantly different from the control (no ComFA). Therefore, the negligible amount of apparent unwinding is likely an artifact of the higher concentrations used. Some protein did apparently stay bound with the DNA because, at higher concentrations of ComFA, a low-mobility complex appeared.

### ATP hydrolysis reduces DNA binding

We showed previously that binding of ComFA to DNA does not require ATP hydrolysis because a Walker B (ATP hydrolysis) mutant bound DNA comparably to wild type protein. This finding left the possibility that ATP hydrolysis could either fuel movement along DNA or release from the DNA (19). Thus, we next investigated ComFA binding and release in the presence or absence of nucleotide analogs. We examined the DNA binding ability of ComFA in the apo state or with AMP-PNP (a non-hydrolyzable ATP analog), ADP, or ATP to determine whether ATP hydrolysis impacts DNA binding. We found that, although ComFA still bound DNA in the presence of 5 μM ATP, its DNA binding was more than seven-fold weaker, with a K_d_ of 171 ± 14 nM, compared with 23.2 ± 0.9 nM, the Kd of the apo state (Figure 5A-C). Five μM AMP-PNP also decreased DNA binding, although not as significantly, with a Kd of 108 ± 25 nM, similar to that of 5 μM ADP (100 ± 45 nM; Figure 5A-C). Of note, the addition of 50 μM or 5 mM ADP reversed the effects of adding 5 μM ADP. The K_d_ of ComFA with 50 μM ADP was significantly reduced compared to the apo state (9.4 ± 1.1 nM; Supplemental Figure S3) and appeared to promote a multimeric or aggregated state, as most of the protein and DNA amassed in the wells at concentrations of protein 50 nM and above (Supplemental Figure S3).

**Figure 5.**
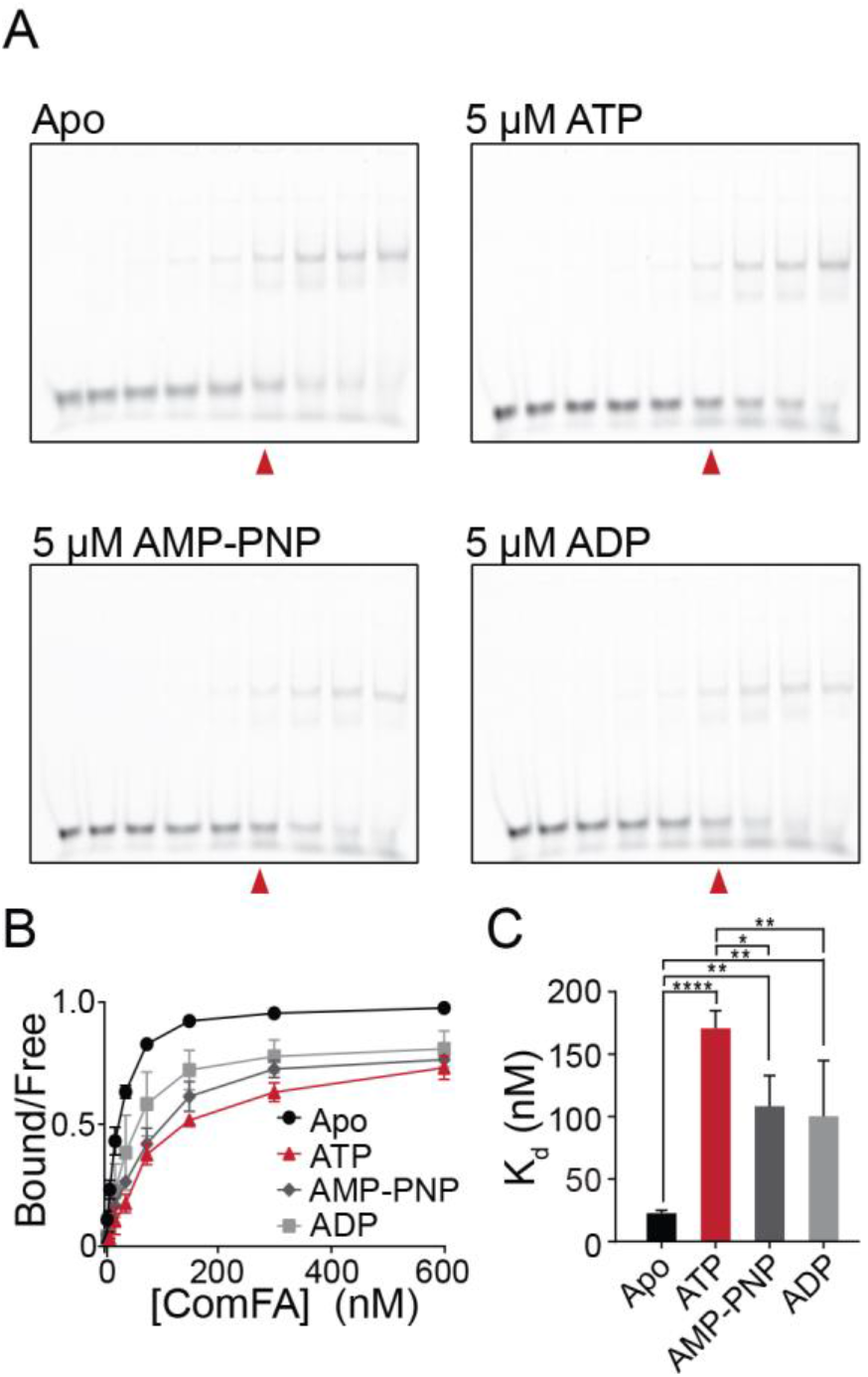
DNA binding of ComFA is weakened in the presence of nucleotide. (A) Representative EMSAs with and without 5 μM ATP, AMP-PNP, or ADP illustrate ComFA binding to 50T DNA substrate (40 nM DNA, 0, 4.8, 9.5, 19, 38, 76, 152, 306, 611 nM ComFA; red triangle indicates 76 nM ComFA on the EMSA). Contrast and brightness were adjusted slightly for presentation. (B-C) Binding curves and mean K_d_ of ComFA in the absence or presence of indicated nucleotide. Apo n=3, ATP n=4, AMP-PNP n=4, ADP n=3. Error bars represent standard deviation. A one-way ANOVA with Sidak’s multiple comparisons test was used for statistical analysis. *p≤0.05, **p≤0.01, ****p≤0.0001

## Discussion

Here, we provide the first experimental evidence that the competence protein, ComFA, translocates on ssDNA in a 5’ to 3’ direction in an ATP-dependent manner. This evidence strongly supports the idea that ComFA acts as a DNA uptake motor in grampositive bacteria and provides a possible mechanism for coupling the ATPase cycle of ComFA to DNA translocation during import.

Our results suggest that ComFA is unusual among DEAD-box helicases, since few DEAD-box helicases interact with DNA, and those with unwinding capabilities primarily melt strands of nucleic acid locally without translocation (28, 31–33) A translocating DEAD-box helicase is not entirely without precedent, since the DEAD-box helicase DDX43 was found to have weak translocase activity on DNA (34). However, the processivity of this protein was limited to fewer than 16 bases (34), making it significantly less processive than ComFA, which we show can travel at least 30-40 bases before DNA release. (34). ComFA also shares homology with PriA, which belongs to the closely-related DExH-box helicases—another member of the helicase superfamily 2. PriA translocates on forked DNA substrates (35,36), and in fact, may play a role in DNA uptake in *Neisseria gonorrhoeae*, a gram-negative bacterium that lacks ComFA (37). Thus, although ComFA contains the DEAD-box motif, functionally it may be more closely related to the translocating DExH family of helicases. Future studies may examine what features give ComFA translocation abilities, setting it apart from other DEAD-box helicases.

Our results that ComFA exhibits a 5’ to 3’ directionality seem to contradict a previous study reporting that DNA uptake occurs in a 3’ to 5’ direction in *S. pneumoniae* (38). These conclusions were based on the authors’ findings that a 3’-labeled strand of DNA was more frequently detected inside the cells than a 5’-labeled strand. One way to reconcile our data with this previous explanation is that ComFA may pull on the degraded strand of DNA from 5’ to 3’, while the transformed strand enters the cell in a 3’ to 5’ manner. Alternatively, this initial report of DNA uptake directionality may have been incorrect in its conclusion that less internalized 5’ label implies a 3’ to 5’ directionality. The protein NucA is thought to cleave one end of the DNA prior to uptake (39), so it is possible that the 5’ end of the DNA is cleaved off prior to uptake in a 5’ to 3’ direction. This would account for the greater quantity of 3’ label detected in the cells. Further, these results have not been replicated in 30 years, and researchers were unable to determine the directionality of uptake in *B. subtilis* using the same method (40). Clearly, the directionality of DNA uptake is a topic that needs further investigation.

We found that ComFA could not unwind dsDNA with the standard helicase substrates we tested, which would suggest that ComFA may act as a motor to drive uptake but likely does not participate directly in unwinding DNA as it enters the cell. This is consistent with the current model of DNA transformation in which strand separation occurs prior to entry into the cell whereupon it would encounter ComFA (1). However, it is possible that ComFA, as with some other helicases, may require a specific conformation of DNA, such as forked DNA, in order to unwind base pairs (41). We have not tested the unwinding ability of ComFA exhaustively with other conformations of DNA, so this leaves open the possibility that ComFA may have some unwinding capability not detected by our assay.

Our finding that ATP hydrolysis causes the release of DNA from ComFA is consistent with the mechanism of other DEAD-box helicases (42). This finding, combined with the discovery that ComFA translocates in a direction-specific manner, allows us to propose a model for the ATPase cycle and DNA interaction, which could aid in understanding the role of ComFA in the DNA uptake complex and guide future experiments. A Brownian ratchet mechanism of translocation seems unlikely, since ComFA exhibits a strong directionality without exogenous forces, such as other competence proteins or a complementary strand of DNA that might force ComFA to move in one direction (43,44). Therefore, we propose that ComFA uses an ‘inchworm’ mechanism for translocation (45,46), utilizing either multiple DNA binding sites on a single ComFA or a ComFA oligomer to move forward (Figure 6). Although we previously reported that ComFA can oligomerize, it is unclear whether the protein is active as a monomer or multimer (19). For the sake of illustration, we assume that ComFA acts as a monomer with two binding sites (BS 1 and BS 2; labeled as 1 and 2 Figure 6), but BS 1 and BS 2 could also represent different ComFA proteins in an oligomer. We demonstrated that ComFA has a higher binding affinity for DNA in its apo state (Figure 5A-D), suggesting that the initial step of DNA binding may be achieved prior to binding nucleotide (Figure 6A). Upon binding to DNA (K_d_ = 23 nM), ComFA binds ATP, leading to a conformational change and higher K_d_ (108 nM; Figure 6B). ATP hydrolysis forces BS 1 to release, yielding a less stable complex (K_d_ = 171 nM; Figure 6C). After ATP hydrolysis, the presence of ADP allows for re-binding of BS 1 and release of BS 2, resulting in the binding state demonstrated by ADP-bound ComFA (K_d_ = 100 nM; Figure 6D). The ADP is then released, and BS 2 rebinds, and the process repeats (Figure 6A). Our finding that a higher concentration of ADP results in protein-DNA aggregates or higher order multimers suggests that ADP or DNA release is the rate-limiting step of the process.

**Figure 6.**
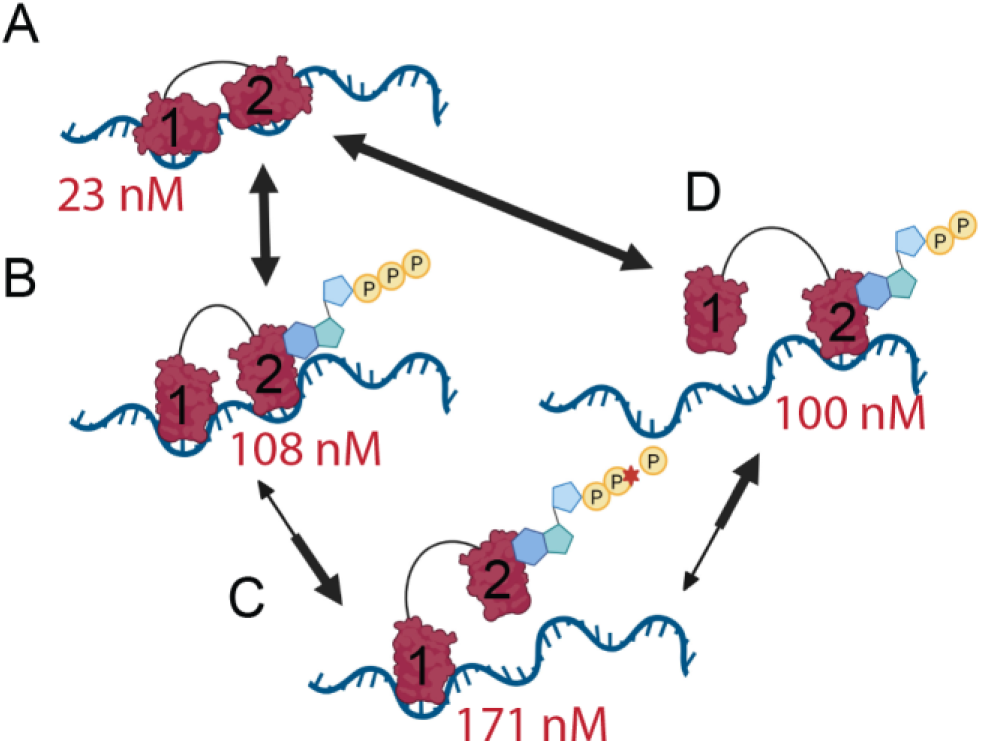
Potential mechanism for ComFA translocation. Maroon proteins labeled 1 and 2 represent either two binding sites on a single ComFA or distinct ComFA proteins in an oligomer bound to ssDNA. (A) ComFA binds ssDNA without nucleotide. (B) ComFA binds ATP, changing conformation leading to (C) ATP hydrolysis and the release of binding site 1. (D) The presence of ADP causes the re-binding of binding site 1 and release of binding site 2. Release of ADP leads to the release of DNA or re-binding of binding site 2 and restoration of original strong binding.

The data we present here strongly support the idea that ComFA may contribute to powering DNA import by translocating along ssDNA, but much remains to be determined. No other DEAD-box helicase has exhibited processive translocation to the extent of ComFA, to our knowledge, and further studies are necessary to confirm the translocase activity of ComFA and examine what sets ComFA apart from other DEAD-box helicases. Further, our data raise questions regarding the directionality of the imported DNA strand. Future studies may also examine the oligomeric state of active ComFA, as well as its behavior within the DNA uptake complex. Protein-protein interactions, particularly with ComEC and ComFC, likely impact multiple parameters of ComFA’s activity, including the speed of translocation and processivity. Finally, we have proposed a model of translocation, but more experiments are necessary to expound on the mechanism of translocation of this unique protein essential to natural transformation in gram-positive bacteria.

## Materials and Methods

### Protein purification

Protein purification was initially completed as described in Diallo et al. (2017;19). Protein purified by this method was used in experiments shown in Figures 1, 2A, 3, as well as Supplemental Figures S1 and S5. Later in the study, we further optimized the protocol including lowering the pH to 6.2 which is more appropriate for purification of the SUMO-ComFA fusion protein. Then, following cleavage of the SUMO tag, we raised the pH for ComFA alone. These preparations of protein were used in Figure 2B-F, 4 and Supplemental Figures S2, S3 and S6. A gel containing purified protein from both the original and revised protocol can be found in Supplemental Figure S4A. Supplemental Figure S4B demonstrates that both protein preps exhibited ATPase activity that was lost in a Walker B (D205A) mutant.

The optimized protocol proceeds as follows, bacteria were grown in LB broth at 42°C to 0.5 OD_600_, followed by induction with a final concentration of 0.5 mM IPTG and incubation at 16°C overnight. Cells were pelleted and resuspended in 3 mL lysis buffer (25 mM Bis-Tris pH6.2, 700 mM NaCl, 5 mM MgCl_2_, 0.13 CaCl_2_, 5 mM imidazole, 1 mM TCEP, 10% glycerol) and then frozen until use. Prior to purification, 3 mL lysis buffer were added to the thawed cells. Cells were disrupted twice using a One-Shot disrupter (Constant Systems). The lysate was then centrifuged for 15 minutes at 20,000 rcf. Supernatant was incubated for 20-40 minutes with 1 mL pre-equilibrated beads (Co-NTA XPure Agarose Resin, pre-charged ion: Cobalt; equilibration buffer: 25 mM Bis-Tris pH6.2, 700 mM NaCl, 5 mM imidazole, 1 mM TCEP, 10% glycerol). Beads were washed (wash buffer: 25 mM Bis-Tris pH6.2, 700 mM NaCl, 20 mM imidazole, 1 mM TCEP, 10% glycerol) and protein was eluted using 4 mL 400 mM imidazole elution buffer (25 mM Bis-Tris pH6.2, 700 mM NaCl, 400 mM imidazole, 1 mM TCEP, 10% glycerol). Protein was then dialyzed overnight (dialysis buffer: 20 mM Bis-Tris pH6.2, 500 mM NaCl, 1 mM TCEP, 10% glycerol) with 100 μl SUMO protease per 1 mL eluate. For proteins to be used in electromobility shift assays (EMSAs), an additional dialysis step (20 mM Bis-Tris pH7.5, 500 mM NaCl, 1 mM TCEP, 5% glycerol) was used to increase the pH.

### Electrophoretic Mobility Shift Assay A (Figure 4)

ComFA at 1.09 - 560 nM was incubated with 40 nM Cy5-labeled oligos, acquired from IDT, in 50 mM Tris-Cl pH 7.5, 50 mM NaCl, 1 mM DTT, 0.1 mg/mL BSA, and 5 mM MgCl_2_ in 25 μL final volume for 30 minutes at room temperature. 3.3% glycerol was added to samples, and 5 μL of each sample was loaded onto a 5% acrylamide 1.5-mm gel in TBE buffer. Gels were pre-run at 75 V for 20 minutes before loading protein/DNA complexes and running at 75 V for 1 hour at 4 °C in 1x TBE running buffer. Gels were imaged on the Azure c600 (Azure Biosystems). For analysis, ImageJ was used to measure integrated density of folded and unfolded DNA.

### Electrophoretic Mobility Shift Assay B (Figures 5, S3, and S4)

Cy5-labeled DNA was acquired from IDT. Protein was diluted into EMSA buffer (2.5 mM CaCl_2_, 12.5 mM MgCl_2_, 75 mM NaCl, 10 mM Tris pH7.5, and 12.5% glycerol) with 0.1mg/mL BSA and 5 nM labeled DNA in a total volume of 20 μl. Samples were incubated for 8-10 minutes, and 16 μl was loaded onto a 5% acrylamide 1.5 mm gel in Tris-Borate-EDTA (TBE) buffer. Gel was pre-run at 85 V for 20 minutes then protein/DNA was added, and the gel was run at 100 V in 0.5x TBE for 1 hour and imaged using a GE Healthcare Typhoon FLA 9000. dsDNA for Figure 1A was created by combining in equal parts 50MB and 50MB-rev in STE buffer (200 mM Tris-HCl, 100 mM EDTA) at a concentration of 1 mM. This mixture was incubated for 5 minutes at 95°C then slowly cooled to room temperature in the heating block over the course of an hour. For analysis, ImageJ was used to measure integrated density bound DNA and free DNA in order to calculate the proportion of bound DNA. The dissociation constant (K_d_) was considered that concentration at which 50% of the DNA was bound.

### ATPase activity assay

ATPase activity was measured using an NADH-coupled plate assay (25) as described in Diallo et al (2017;19). Un-labeled and labeled oligos were obtained from IDT or Sigma Aldrich. All labeled oligos were 50-polyT (50T) and labeled on either end as indicated and were HPLC-purified by the supplier. 20MB30T and 30T20MB were used to demonstrate directionality of ComFA (Table 1). Note that ATPase activity appears lower in Figure 1B than in future experiments because this experiment was completed early in our work on ComFA when we were using higher concentrations of protein. We learned later that high concentrations of the protein actually results in a lower ATPase rate per ComFA (Supplemental Figure S6).

### Modeling

Kinetic models have been developed for ATPases that bind and translocate along DNA that differ based on the underlying mechanism. A set of models that consider a segment of DNA as a lattice of binding sites that allows the ATPase to bind to the lattice and subsequently translocate across the lattice and fall off with different rate constants were proposed by Young et al. (26,27). Below are the two kinetic models presented by Young et al and tested in this work that depend on the length and concentration of the DNA substrate (Scheme II and Scheme III). We compare the fit of the data to these models relative to a null kinetic model (Scheme I) of a non-translocating ATPase that depends only on the concentration of DNA, and not on the length of the lattice. We use an F-test to perform a statistical comparison of these models.

Scheme I:

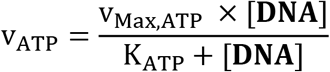

where, v_Max,ATP_ is the saturating ATPase rate and K_ATP_ is the [**DNA**] half-maximal

ATPase rate. This is the null model that is independent of DNA length (**DNALength**).

Scheme II: Scheme I with K_ATP_ being dependent on DNA length.

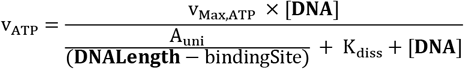

where, v_Max,ATP_ is the saturating ATPase rate, K_diss_ is the dissociation constant of ATPase from DNA and A_uni_ is a composite parameter describing ATPase binding to DNA. bindingSite is the number of nucleotides required for DNA binding by the protein, and (**DNALength**-bindingSite) is the number of unique binding sites available on the DNA.

Scheme III: Scheme I with v_Max,ATP_ being dependent on DNA length.

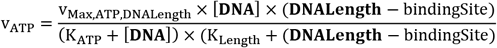

where, v_Max,ATP,DNALength_ is the saturating ATPase rate, K_ATP_ and K_Length_ are composite parameters that depend on the dissociation constant and translocation rate of ATPase on the DNA lattice.

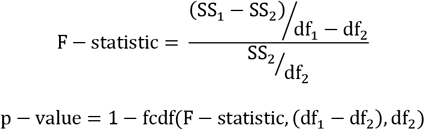

which can be used in hypothesis testing of an alternate model (2) against the null model (1), given that model (1) is a nested model of model (2). SS_x_ represents the sum-squared error of data given a model with df_x_ representing the degrees-of-freedom of a model (number of unique data points – number of fitted parameters). The p-value is estimated as the probability that the null model cannot be discarded given the F-statistic.

Modeling fitting and statistical testing was performed in MATLAB (MATLAB 2018b, The MathWorks, Inc., Natick, Massachusetts, United States.) using th

## Data Availability

MATLAB code and raw data are made available at (https://github.com/TheBurtonLab/Foster-et-al.).

## Strand Displacement Assay

Duplex DNA (dsDNA) structures 3’ ds long/20MB-Cy5 or 30T20MB/20MB-Cy5 were annealed in 10 nM Tris-Cl pH 7.5 and 100 mM NaCl to a final concentration of 5 μM by heating to 95 **°**C in a heat block and slowly cooling to room temperature over several hours. ComFA at 1.09 - 560 nM was incubated with 40 nM oHF172/oHF174-Cy5 or oHF169/oHF175-Cy5 in 50 mM Tris-Cl pH 7.5, 50 mM NaCl, 1 mM DTT, 0.1 mg/mL BSA, 5 mM MgCl_2_, 4 mM ATP, and 4 μM trap oligo (unlabeled 20MB30T) in 25 μL final volume for 15 minutes at 37 °C. Melted controls were heated to 95 °C for 10 minutes. Reactions were quenched using 5 μL stop buffer (2% Sodium dodecyl sulfate, 5 μg/mL Proteinase K, 20% glycerol, and 0.1 M EDTA), and 5 μL of each sample were loaded on 15% acrylamide 1.5-mm gels in Tris-Borate-EDTA (TBE) buffer. Gels were run for 1 hour at 75 V at 4 °C in 1x TBE running buffer. Gels were imaged using the Azure c600 (Azure Biosystems).

## Acknowledgements

We would like to thank Drs. Michael M. Cox, Aaron A. Hoskins, and Katarzyna A. Gromek for helpful discussions and feedback.

## References

1. Burton, B. and Dubnau, D. (2010) Membrane-associated DNA Transport Machines. Cold Spring Harb. Perspect. Biol., 2, a000406–a000406.

2. Chen, I., Christie, P.J. and Dubnau, D. (2005) The Ins and Outs of DNA Transfer in Bacteria. Science, 310, 1456–1460.

3. Johnston, C., Martin, B., Fichant, G., Polard, P. and Claverys, J.-P. (2014) Bacterial transformation: distribution, shared mechanisms and divergent control. Nat. Rev. Microbiol., 12, 181–196.

4. Johnsborg, O., Eldholm, V. and Håvarstein, L.S. (2007) Natural genetic transformation: prevalence, mechanisms and function. Res. Microbiol., 158, 767–778.

5. Lorenz, M.G. and Wackernagel, W. (1994) Bacterial gene transfer by natural genetic transformation in the environment. Microbiol. Rev., 10.1128/mmbr.58.3.563-602.1994.

6. Maier, B., Chen, I., Dubnau, D. and Sheetz, M.P. (2004) DNA transport into Bacillus subtilis requires proton motive force to generate large molecular forces. Nat. Struct. Mol. Biol., 11.

7. Inamine, G.S. and Dubnau, D. (1995) ComEA, a Bacillus subtilis integral membrane protein required for genetic transformation, is needed for both DNA binding and transport. J. Bacteriol., 177, 3045–3051.

8. Provvedi, R. and Dubnau, D. (1999) ComEA is a DNA receptor for transformation of competent Bacillus subtilis. Mol. Microbiol., 31, 271–280.

9. Dubnau, D. and Cirigliano, C. (1972) Fate of transforming deoxyribonucleic acid after uptake by competent Bacillus subtilis: size and distribution of the integrated donor segments. J. Bacteriol., 111, 488–494.

10. Davidoff-Abelson, R. and Dubnau, D. (1973) Conditions affecting the isolation from transformed cells of Bacillus subtilis of high-molecular-weight singlestranded deoxyribonucleic acid of donor origin. J. Bacteriol., 116, 146–153.

11. Piechowska, M. and Fox, M.S. (1971) Fate of Transforming Deoxyribonucleate in Bacillus subtilis. J. Bacteriol., 108, 680–689.

12. Hahn, J., Albano, M. and Dubnau, D. (1987) Isolation and characterization of Tn917lac-generated competence mutants of Bacillus subtilis. J. Bacteriol., 169, 3104–3109.

13. Draskovic, I. and Dubnau, D. (2005) Biogenesis of a putative channel protein, ComEC, required for DNA uptake: membrane topology, oligomerization and formation of disulphide bonds. Mol. Microbiol., 55, 881–896.

14. Levine, J.S. and Strauss, N. (1965) LAG PERIOD CHARACTERIZING THE ENTRY OF TRANSFORMING DEOXYRIBONUCLEIC ACID INTO BACILLUS SUBTILIS. J. Bacteriol., 89, 281–287.

15. Strauss, N. (1965) CONFIGURATION OF TRANSFORMING DEOXYRIBONUCLEIC ACID DURING ENTRY INTO BACILLUS SUBTILIS. J. Bacteriol., 89, 288–293.

16. Londoño-Vallejo, J.A. and Dubnau, D. (1994) Membrane association and role in DNA uptake of the Bacillus subtilis PriA analogue ComF1. Mol. Microbiol., 13, 197–205.

17. Londoño-Vallejo, J.A. and Dubnau, D. (1994) Mutation of the putative nucleotide binding site of the Bacillus subtilis membrane protein ComFA abolishes the uptake of DNA during transformation. J. Bacteriol., 176, 4642–4645.

18. Chilton, S.S., Falbel, T.G., Hromada, S. and Burton, B.M. (2017) A Conserved Metal Binding Motif in the Bacillus subtilis Competence Protein ComFA Enhances Transformation. J. Bacteriol., 199.

19. Diallo, A., Foster, H.R., Gromek, K.A., Perry, T.N., Dujeancourt, A., Krasteva, P.V., Gubellini, F., Falbel, T.G., Burton, B.M. and Fronzes, R. (2017) Bacterial transformation: ComFA is a DNA-dependent ATPase that forms complexes with ComFC and DprA. Mol. Microbiol., 105, 741–754.

20. Lee, M.S., Dougherty, B.A., Madeo, A.C. and Morrison, D.A. (1999) Construction and analysis of a library for random insertional mutagenesis in Streptococcus pneumoniae: use for recovery of mutants defective in genetic transformation and for identification of essential genes. Appl. Environ. Microbiol., 65, 1883–1890.

21. Bergé, M., Moscoso, M., Prudhomme, M., Martin, B. and Claverys, J.-P. (2002) Uptake of transforming DNA in Gram-positive bacteria: a view from Streptococcus pneumoniae. Mol. Microbiol., 45, 411–421.

22. Andersen, C.B.F., Ballut, L., Johansen, J.S., Chamieh, H., Nielsen, K.H., Oliveira, C.L.P., Pedersen, J.S., Séraphin, B., Hir, H.L. and Andersen, G.R. (2006) Structure of the Exon Junction Core Complex with a Trapped DEAD-Box ATPase Bound to RNA. Science, 313, 1968–1972.

23. Hilbert, M., Karow, A.R. and Klostermeier, D. (2009) The mechanism of ATP-dependent RNA unwinding by DEAD box proteins. Biol. Chem., 390.

24. Kikuma, T., Ohtsu, M., Utsugi, T., Koga, S., Okuhara, K., Eki, T., Fujimori, F. and Murakami, Y. (2004) Dbp9p, a member of the DEAD box protein family, exhibits DNA helicase activity. J. Biol. Chem., 279, 20692–20698.

25. Gilbert, S.P. and Mackey, A.T. (2000) Kinetics: A Tool to Study Molecular Motors. Methods, 22, 337–354.

26. Young, M.C., Kuhl, S.B. and von Hippel, P.H. (1994) Kinetic Theory of ATP-driven Translocases on One-dimensional Polymer Lattices. J. Mol. Biol., 235, 1436–1446.

27. Young, M.C., Schultz, D.E., Ring, D. and von Hippel, P.H. (1994) Kinetic Parameters of the Translocation of Bacteriophage T4 Gene 41 Protein Helicase on Singlestranded DNA. J. Mol. Biol., 235, 1447–1458.

28. Linder, P. and Jankowsky, E. (2011) From unwinding to clamping — the DEAD box RNA helicase family. Nat. Rev. Mol. Cell Biol., 12, 505–516.

29. Fischer, C.J. and Lohman, T.M. (2004) ATP-dependent translocation of proteins along single-stranded DNA: models and methods of analysis of pre-steady state kinetics. J. Mol. Biol., 344, 1265–1286.

30. Nelson, S.W., Perumal, S.K. and Benkovic, S.J. (2009) Processive and Unidirectional Translocation of Monomeric UvsW Helicase on Single-Stranded DNA†. 10.1021/bi801792q.

31. Putnam, A.A. and Jankowsky, E. (2013) DEAD-box helicases as integrators of RNA, nucleotide and protein binding. Biochim. Biophys. Acta BBA - Gene Regul. Mech., 1829, 884–893.

32. Yang, Q., Del Campo, M., Lambowitz, A.M. and Jankowsky, E. (2007) DEAD-Box Proteins Unwind Duplexes by Local Strand Separation. Mol. Cell, 28, 253–263.

33. Gilman, B., Tijerina, P. and Russell, R. (2017) Distinct RNA-unwinding mechanisms of DEAD-box and DEAH-box RNA helicase proteins in remodeling structured RNAs and RNPs. Biochem. Soc. Trans., 45, 1313–1321.

34. Talwar, T., Vidhyasagar, V., Qing, J., Guo, M., Kariem, A., Lu, Y., Singh, R.S., Lukong, K.E. and Wu, Y. (2017) The DEAD-box protein DDX43 (HAGE) is a dual RNA-DNA helicase and has a K-homology domain required for full nucleic acid unwinding activity. J. Biol. Chem., 292, 10429–10443.

35. Sandler, S.J. and Marians, KJ. (2000) Role of PriA in Replication Fork Reactivation in Escherichia coli. J. Bacteriol., 182, 9–13.

36. Lasken, R.S. and Kornberg, A. (1988) The primosomal protein n’ of Escherichia coli is a DNA helicase. J. Biol. Chem., 263, 5512–5518.

37. Kline, K.A. and Seifert, H.S. (2005) Mutation of the priA Gene of Neisseria gonorrhoeae Affects DNA Transformation and DNA Repair. J. Bacteriol., 187, 5347–5355.

38. Vincent Mejean and Claverys, J.-P. (1988) Polarity of DNA entry in transformation of Streptococcus pneumoniae. Mol. Gen. Genet. 213(2-3):444–8

39. Provvedi, R., Chen, I. and Dubnau, D. (2001) NucA is required for DNA cleavage during transformation of Bacillus subtilis. Mol. Microbiol., 40, 634–644.

40. Vagner, V., Claverys, J.P., Ehrlich, S.D. and Méjean, V. (1990) Direction of DNA entry in competent cells of Bacillus subtilis. Mol. Microbiol., 4, 1785–1788.

41. Brosch, R.M. and Matson, S.W. (2020) History of DNA Helicases. Genes, 11, 255.

42. Liu, F., Putnam, A. and Jankowsky, E. (2008) ATP hydrolysis is required for DEAD box protein recycling but not for duplex unwinding. Proc. Natl. Acad. Sci., 105, 20209–20214.

43. Donmez, I. and Patel, S.S. (2006) Mechanisms of a ring shaped helicase. Nucleic Acids Res., 34, 4216–4224.

44. Levin, M.K., Gurjar, M. and Patel, S.S. (2005) A Brownian motor mechanism of translocation and strand separation by hepatitis C virus helicase. Nat. Struct. Mol. Biol., 12, 429–435.

44. Mackintosh, S.G. and Raney, K.D. (2006) DNA unwinding and protein displacement by superfamily 1 and superfamily 2 helicases. Nucleic Acids Res., 34, 4154–4159.

45. Patel, S.S. and Donmez, I. (2006) Mechanisms of Helicases. J. Biol. Chem., 281, 18265–18268.

